# The SNARC effect involves both representation and response-selection processing stages: Evidence from ERPs

**DOI:** 10.1101/2023.11.01.565045

**Authors:** Yang Weibin, Lizhu Yan, Xinrui Xiang, Weizhi Nan

**Affiliations:** Department of Psychology and Center for Brain and Cognitive Sciences, School of Education, Guangzhou University, Guangzhou 510006, China

**Keywords:** SNARC, two-stage processing model, P1, N2, P300

## Abstract

The processing stage (i.e., the early semantic representation stage, the late response-selection stage, or both) at which the spatial-numerical association of response codes (SNARC) effect occurs is still controversial. The two-stage processing model hypothesizes that the SNARC effect involves both stages and that different interference factors acting at the two stages might be the core reason for the observed stage flexibility of the SNARC effect. To test this hypothesis, the present study was designed to elicit the SNARC, Stroop (semantic-representation stage related), and Simon (response-selection stage related) effects together in one magnitude comparison task and used the event-related potentials (ERPs) to observe the temporal dynamics of these effects. The behavioral results showed no interaction between the Stroop and Simon effects, while these two effects both interacted with the SNARC effect. Furthermore, the ERP results showed an interaction of the Stroop effect with the SNARC effect for the early sensory P1 component, while the interaction of the Simon effect with the SNARC effect was evident for the late N2 and P300 components. The current study repeatedly verified the independence of Stroop and Simon effects. Most importantly, the temporal-specific interactions among the SNARC effect and the other two stage-related factors provided further evidence to support the two-stage processing model that the SNARC effect involves both the representation and response-selection stages.

## Introduction

The spatial-numerical association of response codes effect (SNARC) refers to the phenomenon in which the response latency for the left effector is shorter than that for the right effector for small numbers (e.g., 1 or 2) and vice versa for larger numbers (e.g., 8 or 9) (Dehaene et al., 1993). According to the mental number line account, numbers are internally represented on a left-to-right mental number axis (Bächtold et al., 1998; Dehaene et al., 1993). The semantic representation of number magnitude (smaller or larger) automatically activates the associated spatial representation on the mental number axis, with smaller numbers mapping onto the left side and larger number mapping onto the right side (Fias, 1996; Fischer et al., 2004; Lammertyn et al., 2002). When the activated spatial representation is congruent with the assigned response effectors, responses are faster and more accurate than when they are incongruent. Previous studies have confirmed the stability of the SNARC effect through various experimental manipulations, such as by expanding the number range to double-digit numbers (Brysbaert, 1995) and negative numbers (Kong et al., 2012), expanding the stimuli to musical notes (Prpic et al., 2016), pitch stimuli (Weis et al., 2016), and luminance stimuli (Fumarola et al., 2014), and expanding the type of response effectors to hand crossing (Dehaene et al., 1993) and foot responses (Hartmann et al., 2014).

However, there is still controversy regarding its underlying mechanism about which processing stage the SNARC effect occurs in (Yan et al., 2022). Researchers have considered the following three perspectives in determining the processing stage in which the SNARC effect occurs: (1) examining the relationship between the SNARC effect and other effects (e.g., the Simon effect, Stroop effect, numerical distance effect, and switching costs) occurring in different stages (Keus & Schwarz, 2005; Mapelli et al., 2003), (2) examining changes in the SNARC effect with different response effectors (e.g., hand responses, eye movements) (Dehaene et al., 1993; Pinto et al., 2019), and (3) analyzing the components of the event-related potentials (ERPs) induced by the SNARC effect (Gut et al., 2012; Keus et al., 2005). Three contradictory views have been proposed: the SNARC effect occurs only in the semantic representation stage (Fischer et al., 2004; Gut et al., 2012; Mapelli et al., 2003; Pinto et al., 2019), only in the response selection stage (Gevers et al., 2005; Keus & Schwarz, 2005; Yan et al., 2021), or flexibly in both stages (Moro et al., 2018; Nan et al., 2022).

First, exploring the relationship between the SNARC effects and other effects occurring in a specific stage is a practicable approach to locate the SNARC effect (Keus & Schwarz, 2005; Mapelli et al., 2003; Yan et al., 2021). Notably, the additive factor method (AFM) suggests that an interaction of two effects indicates that both effects occur in a common processing stage (Sternberg, 1969). For instance, it has been found that the Stroop effect occurs in the early semantic representation stage (Hock & Egeth, 1970; Li et al., 2014), while the Simon effect occurs in the late response selection stage (Ansorge, 2003; De Jong et al., 1994; Leuthold, 2011). Thus, if either of these two effects interacts with the SNARC effect, the observed interaction could help to verify at which stage the SNARC effect occurs. Mapelli et al. (2003) simultaneously elicited SNARC and Simon effects in one task and found that two effects were additive, which implied that the SNARC effect might not occur in the response selection stage alongside the Simon effect, rather occurring in the early semantic representation stage (see also in Tlauka, 2002). Yan et al. (2021) further simultaneously elicited SNARC, Simon, and Stroop effects and found that the SNARC effect interacted only with the Simon effect and was independent of the Stroop effect (see also in Gevers et al., 2005), consistent with the results of Mapelli et al. (2003) showing that the SNARC effect occurs in the early semantic representation stage. However, Nan et al. (2022) applied a similar paradigm to that used in Yan et al. (2021) and found that the SNARC effect interacted with both the Stroop and Simon effects, implying that the SNARC effect might occur in both the semantic representation and response selection stages. Using the logic of the AFM, some researchers considered other effects (e.g., the distance effect and switching costs in Moro et al., 2018; the switching costs in Zhang et al., 2021) and found inconsistent interactions with the SNARC effect. In general, a unanimous conclusion about the processing stage of the SNARC effect has not been drawn from the evidence obtained given this perspective.

Second, another way to verify the processing stage of the SNARC effect is to manipulate what is relative to certain processing stages and monitor the changes in the SNARC effect. Fischer et al. (2004) observed the SNARC effect in eye movement where no lateral response codes were involved, which indicates that the SNARC effect occurs in the early semantic representation stage. Pinto et al. (2019) applied a unimanual go/no-go paradigm that does not require the use of lateral response codes to explore the contribution of the codes of the semantic representation stage. Their results indicated that the SNARC effect was observed only with the joint activation of contrasting semantic-related spatial codes and number magnitude codes, which also indicates that the SNARC effect occurs in the early semantic representation stage. However, Keus and Schwarz (2005) manipulated the subjects’ response effector (vocal or manual) in their experiments and found that executing a lateral response (manual response) is essential for the SNARC effect, which indicates that the SNARC effect occurs in the late response selection stage. Together, the evidence reflecting the processing stage of the SNARC effect obtained from this perspective is also still inconclusive.

Third, a direct way to observe the temporal dynamics of the occurrence of the SNARC effect is to use electrophysiological measurements. Some ERP studies found that the early N1 and P1 components were sensitive to the SNARC effect, which supports the involvement of the early semantic representation stage (Gut et al., 2012; Schuller et al., 2015). However, Keus et al. (2005) found that the lateralized readiness potential (LRP) component, which is related to response preparation and execution, was also sensitive to the SNARC effect, which supports the involvement of the response selection stage (see also in Gevers et al., 2006).

Overall, the processing stage in which the SNARC effect occurs is still controversial. Conflicting evidence may be caused by the following two factors: (1) disparities in the comprehension of additive-factor logic leading to indirect inference; and (2) observations from a single point leading to indirect inference (Xiang et al., 2022; Yan et al., 2022). According to the assumptions of the AFM, the only inference we can draw from a noninteraction observation is heterogeneity, rather than a precise localization of the processing stage. Therefore, the AFM results in an indirect inference that treats the proposition that the SNARC effect occurs in the early semantic representation stage as a corollary of the observed noninteraction between the SNARC and Simon effects (Mapelli et al., 2003; Tlauka, 2002). A direct inference requires the comprehensive observation of the relationship among the SNARC effect, an effect related to the semantic representation stage, and an effect related to the response selection stage. However, conclusive evidence has not been obtained even with such a comprehensive observation. Although studies by Nan et al., (2022) and Yan et al., (2021) have evoked the SNARC, Stroop, and Simon effects simultaneously, contradictory results have been reported. This contradiction may arise from the use of different types of Stroop effects, in which conflict arises from parity information vs. magnitude information (Xiang et al., 2022). Notably, ERPs with their high temporal resolution can be deemed another source of direct evidence. However, there is a lack of observations of the temporal dynamics of the relationship among the SNARC effect and other effects related to a particular stage. Both the studies by Keus et al. (2005) and by Gevers et al. (2006) found that the P300 component was sensitive to the SNARC effect, which supports the involvement of the response selection stage. Nonetheless, it cannot be ruled out that the SNARC effect may also involve the early semantic representation stage.

Accounting for the two reasons mentioned above, based on the study of Nan et al. (2022), the present study aimed to comprehensively observe the SNARC, Stroop, and Simon effects in one magnitude comparison task by using the ERP technique. With the high temporal resolution characteristics of EEG, we can directly observe the temporal dynamics of these three effects and their interactions. Since the SNARC effect results from a conflict within the domain of number recognition, it may involve both the encoding of the number representation and the cognitive resolution of the conflict. Traditionally, the P1 component is regarded as the earliest endogenous visual ERP component linked to the sensory analysis of stimulus features (Paz-Caballero & García-Austt, 1992; Taylor, 2002). N2 and P300 components are sensitive to conflict monitoring and resolution, respectively (Groom & Cragg, 2015; Li et al., 2015; Wang et al., 2014). Thus, the interactions among the three effects on the sensory P1, N2, and P300 components were monitored. Three hypotheses were proposed: (1) the behavioral result from Nan et al. (2022) would be replicated in that the SNARC effect would interact with the Stroop and Simon effects; (2) the SNARC effect would show an interaction with the magnitude Stroop effect at the early ERP component because of the involvement of the early semantic representation stage; and (3) the interaction between the Simon and SNARC effects would be observed at the late components because of the involvement of the late response selection stage.

## Materials and Methods

### Participants

The required sample size was estimated by G*power 3.1 (Faul et al., 2007). Given a conservative estimate of statistical power for detecting an effect at an alpha level of 0.05, a minimum sample of 23 participants was deemed necessary.

Twenty-six undergraduate students (11 males, age range: 18–25 years old) participated. All the participants reported having no neurological or psychiatric history; they were right-handed and had normal or corrected-to-normal vision. Each participant voluntarily enrolled and signed an informed consent form before the experiments. The Institutional Review Board of the Educational School, Guangzhou University approved this study.

### Stimuli, design, and procedure

The participants were seated in a sound-attenuating chamber approximately 43 cm away from a liquid-crystal display (LCD) monitor (resolution 1024 × 768 pixels, vertical refresh rate 75 Hz), with their eyes at the same level as the center of the monitor. All stimuli were displayed on a gray panel (300 × 300 pixels) on a black background. Stimulus presentation and response recording were controlled by E-Prime 3.0 software (Psychological Software Tools, Inc., Pittsburgh, PA).

The stimuli were presented in the center of the screen; these stimuli consisted of numbers and Chinese characters, with each number overlaid on a character. Specifically, the stimuli varied by the numbers (1, 2, 3, 4, 6, 7, 8, 9; presented in boldface with a size of 1 × 1.5°), the rotation of the numbers (clockwise rotation with 20° or -20° with 0° as 12 o’clock), and the background text (Chinese characters: “大” or “小”, meaning “large” and “small,”, respectively; presented in a regular script in red with a size 5 × 6°).

A task with a 2 (SNARC effect: congruent and incongruent) × 2 (Stroop effect: congruent and incongruent) × 2 (Simon effect: congruent and incongruent) within-subject design was performed. The SNARC effect was manipulated by the response rule (congruent: small/large numbers with the left/right-hand response, respectively; incongruent: small/large numbers with the right/left-hand response, respectively); the Simon effect was manipulated by the number orientation and response hand (congruent: number rotation with -20°/20° with left/right-hand response, respectively; incongruent: number rotation with -20°/20° with right/left-hand response, respectively); and the Stroop effect was manipulated by the number magnitude and the background character (congruent: large/small numbers with the “大”/“小” character, respectively; incongruent: large/small numbers with the “小”/“大” character, respectively). According to the difference among the stimuli presented, the trials were classified into eight conditions, called “SNCStCSiC”, “SNCStCSiI”, “SNCStISiC”, “SNCStISiI”, “SNIStCSiC”, “SNIStCSiI”, “SNIStISiC” and “SNIStISiI”. In this notation, “SN” refers to the SNARC effect, “St” to the Stroop effect, and “Si” to the Simon effect; “C” indicates congruence, and “I” indicates incongruence; for example, trials classified as the “SNCStCSiI” condition have a congruent SNARC effect and Stroop effect, while the Simon effect is incongruent.

The formal experiment consisted of eight blocks; each block consisted of 80 trials, resulting in a total of 640 trials, with 80 trials for each condition. Four blocks were used for SNARC congruent response mapping (small/large numbers with the left/right-hand response, respectively). The other four blocks were used for SNARC incongruent response mapping (small/large numbers with the right/left-hand response, respectively). The stimulus-response mapping order was counterbalanced among subjects.

There were four practice blocks before the formal test. Practice block 1 consisted of 50 trials in which the stimuli were numbers (1–9, except 5) or Chinese characters (“大” and “小”) to strengthen the connection between numbers and Chinese characters for the Stroop effect. Practice block 2 consisted of 50 trials using stimuli of the letters A and B (rotated at -20°/20°) to strengthen the connection between the stimulus rotation and the response hand for the Simon effect. Practice block 3 consisted of 40 trials and used the numbers 3 and 6 (rotated at with -20°/20°) as stimuli to test the Simon effect. If there was no evidence of the Simon effect, practice block 2 would be presented again. Practice block 4 consisted of 10 trials that were similar to those in the formal experiment to familiarize the participants with the formal experimental.

As shown in Figure 1, each trial began with the presentation of a fixation cross (with a size of 0.7 × 0.7°) for 500 ms, followed by the target with a black number (clockwise rotation with -20°/+20°) superimposed on a red Chinese character. The target was presented until a response was given or 1000 ms had elapsed. Participants were instructed to respond to the number by pressing a button as rapidly and accurately as possible (the “S” button for small numbers with the left index finger and the “L” button for large numbers with the right index finger in the SNARC congruent condition) while ignoring its rotation and the background Chinese character. The target disappeared when participants pressed a button, and then a fixation cross was presented in the center of the screen for 500–1000 ms.

**Figure 1.**
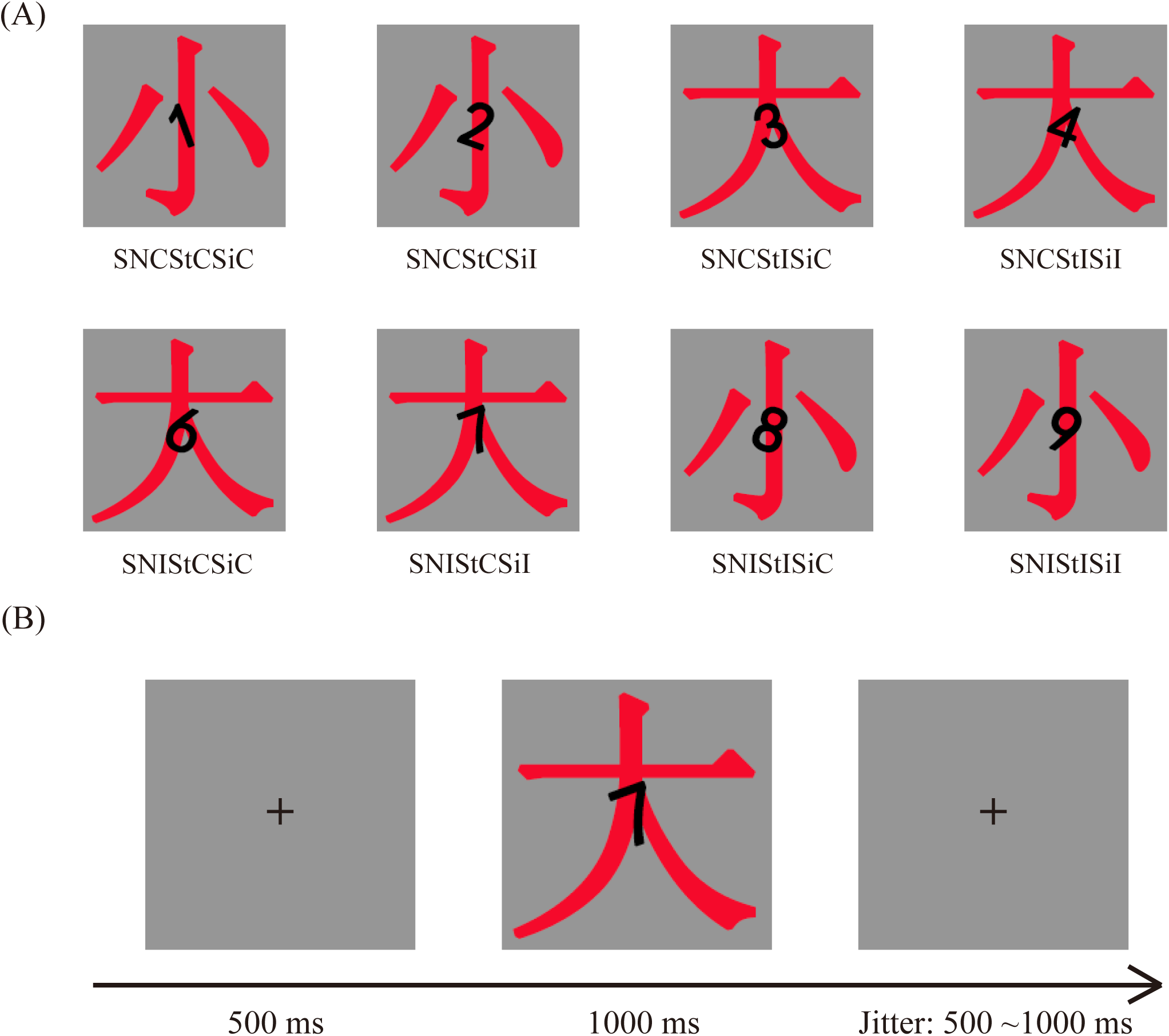
A) Conditions for the magnitude comparison task eliciting the Stroop, Simon, and/or SNARC effects. There are eight conditions in total, each with three effect attributes. The naming conventions of the eight conditions are as follows: “SN” represents SNARC effect, “St” represents Stroop effect, “Si” represents Simon effect, “C” indicates congruence, and “I” indicates incongruence. For example, “SNCStCSiC” indicates that the trial is SNARC congruent, Stroop congruent and Simon congruent. Other trials are named accordingly. Specifically, when using the left hand response rule for a small number, the SNARC effect is congruent. For the Stroop effect, if the background text is “small”, then presentation of the number 3 is considered congruent, while that of the number 9 is incongruent. For the Simon effect, the number 3 presented to the left is considered congruent, while number 3 presented to the right is incongruent. B) Schematic illustration of the magnitude comparison task. At the beginning, a fixation cross is shown for 500 ms, followed by the presentation of the target to which participants are instructed to respond by judging its magnitude. After the target disappears, a fixation cross is presented at the center of the screen. Its presentation time was taken randomly from the range of 500–1000 ms.

**Figure 2.**
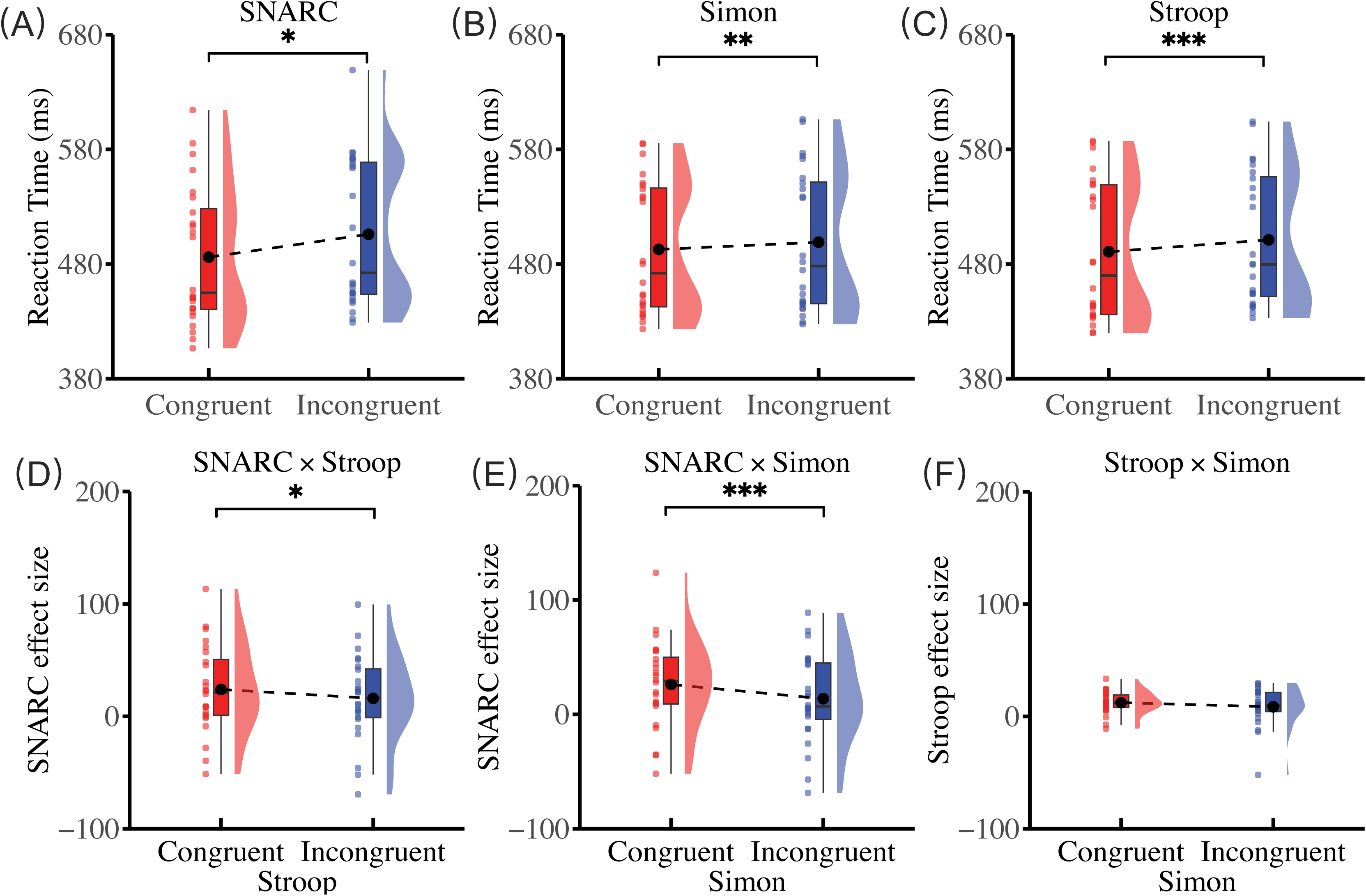
Summary of behavioral results. The upper panels show the main effect of (A) SNARC, (B) Simon and (C) Stroop effects. The lower panels show the interaction results.

**Figure 3.**
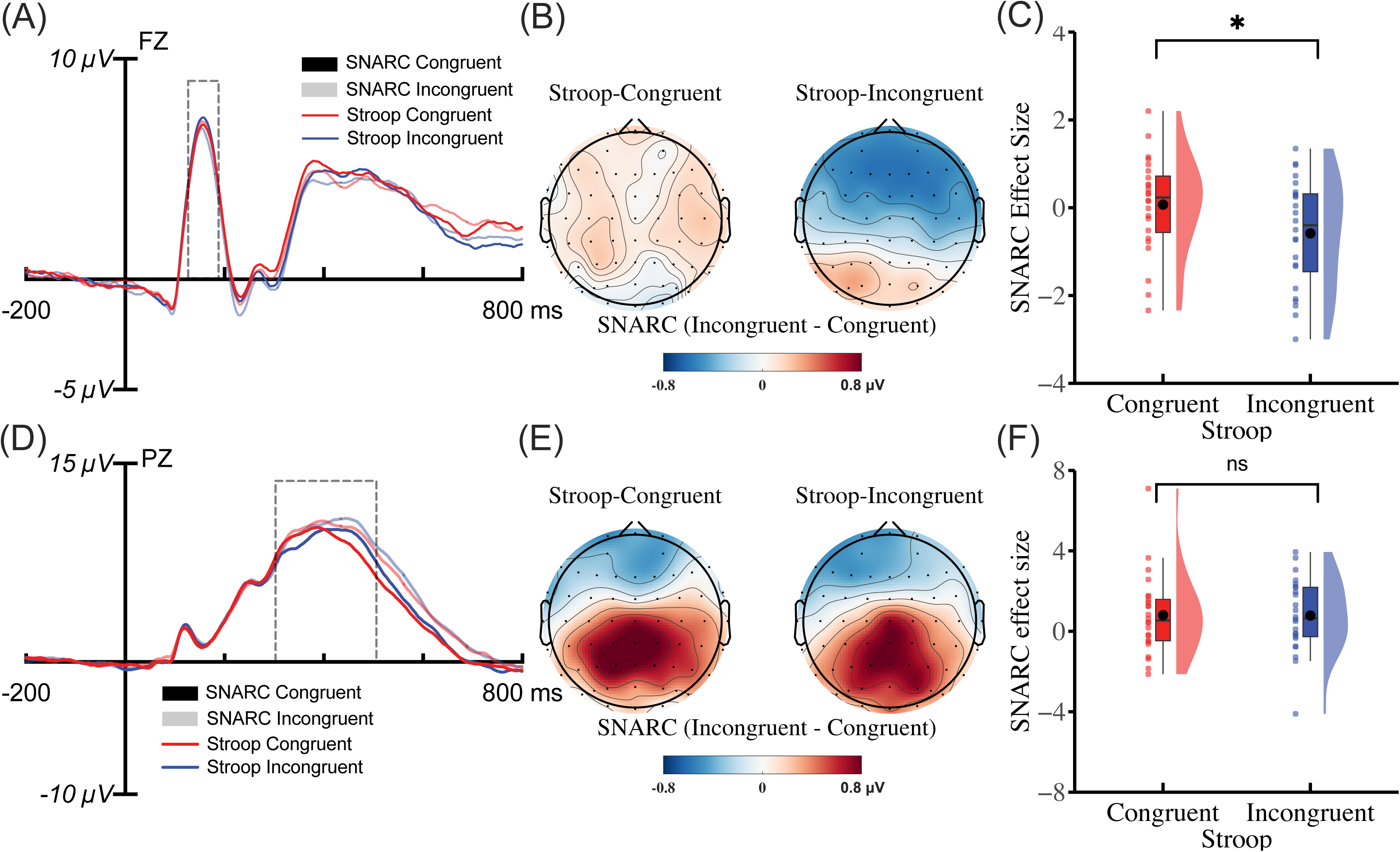
(A) The interaction between the SNARC and Stroop effects for the P1 component. The mappings of the waveform and condition representations are as follows: the color of the waveform represents the congruency of the Stroop effect, and the transparency of the waveform represents the congruency of the SNARC effect. For example, the red translucent waveform corresponds to the SNARC incongruent and Stroop congruent trials, and the blue opaque waveform corresponds to SNARC congruent and Stroop incongruent trials. (B) Topographies were calculated by averaging the data within a time window of 121 to 195 ms after the onset of the target. The topography of the SNARC effect (i.e., the subtraction of amplitudes under congruent and incongruent conditions) in Stroop congruent trials is presented on the left, and that of Stroop incongruent trials is presented on the right. (C) The raincloud plot of the SNARC effect for the P1 component in Stroop congruent and incongruent conditions. (D) The interaction between the SNARC and Stroop effects for the P300 component. (E) Topographies were calculated by averaging the data within a time window of 311 to 497 ms after the onset of the target. (F) The raincloud plot of the SNARC effect on the P300 component in Stroop congruent and incongruent conditions.

**Figure 4.**
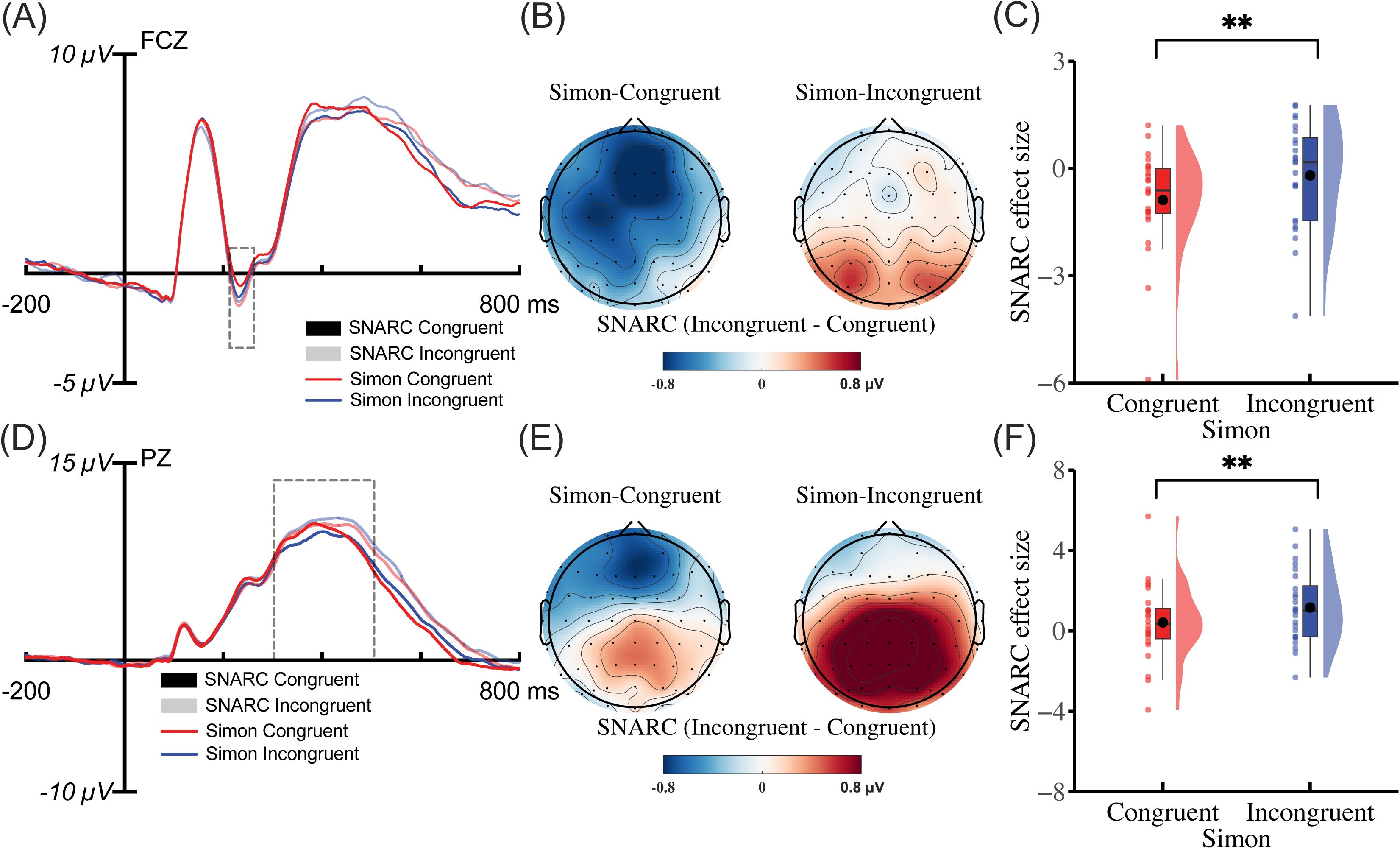
(A) The interaction between the SNARC and Simon effects for the N2 component. The color of the waveform represents the congruency of the Simon effect, and the transparency of the waveform represents the congruency of the SNARC effect. For example, the red translucent waveform corresponds to SNARC incongruent and Simon congruent trials, and the blue opaque waveform corresponds to SNARC congruent and Simon incongruent trials. (B) Topographies of the SNARC effect (i.e., the subtraction of amplitudes under congruent and incongruent conditions) were calculated by averaging the data within a time window of 220 to 245 ms after the onset of the target. (C) The raincloud plot of the SNARC effect for the N2 component in the Simon congruent and incongruent conditions. (D) The interaction between the SNARC and Simon effects for the P300 component. (E) Topographies were calculated by averaging the data within a time window of 311 to 497 ms after the onset of the target. (F) The raincloud plot of the SNARC effect for the P300 component in the Simon congruent and incongruent conditions.

**Figure 5.**
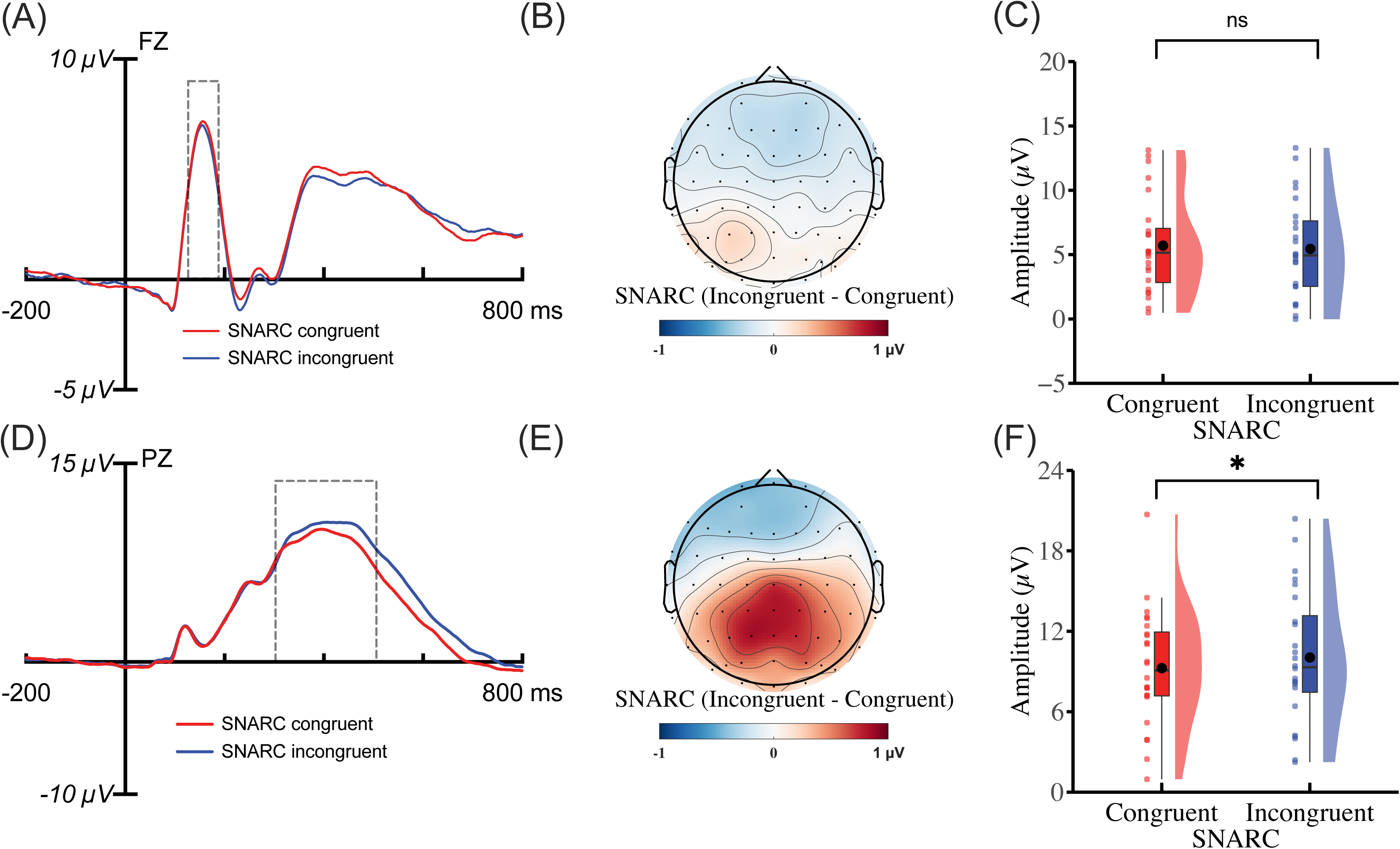
(A) The SNARC effect for the P1 component. The color of the waveform represents the congruency of the SNARC effect. The red waveform corresponds to SNARC congruent trials, and the blue waveform corresponds to SNARC incongruent trials. (B) Topographies of the SNARC effect (i.e., the subtraction of amplitudes under congruent and incongruent conditions) were calculated by averaging the data within a time window of 121 to 195 ms after the onset of the target. (C) Raincloud plot of the SNARC effect on the P1 component. (D) The SNARC effect for the P300 component. (E) Topographies were calculated by averaging the data within a time window of 311 to 497 ms after the onset of the target. (F) The raincloud plot of the SNARC effect for the P300 component.

### EEG Recording and Preprocessing

EEG data were acquired through a 64-electrode Ag/AgCI system (NeuroScan, Texas, USA) in AC mode. Electrodes were positioned according to the 10/20 International System and the electrode between FPz and Fz was used as ground. All electrodes were referenced to the electrode placed between FCz and Cz during the recording and were rereferenced offline to the average of the left and right mastoids. The horizontal and vertical electrooculograms (EOGs) were recorded using electrodes placed at the left and right external canthi and above and below the left eye. EEG and EOG were sampled at 1000 Hz and filtered with a 0.05–100 Hz bandpass filter using a Neuroscan SynAmps digital amplifier system (Neuroscan Labs, Sterling, VA). Electrode impedances were kept below 5 kΩ throughout the experiment.

After the acquisition, all data analyses were performed in MATLAB using the EEGLAB Toolbox (Delorme & Makeig, 2004) and ERPLAB Toolbox (Lopez-Calderon & Luck, 2014). Before artifact rejection and correction, the signals were bandpass filtered using a noncausal IIR Butterworth filter (high amplitude cutoff = 0.05 Hz, low amplitude cutoff = 30 Hz, slope = 12 dB/octave). Subsequently, the data were segmented into epochs of 1000 ms in length, including a 200 ms prestimulus baseline interval for baseline correction.

For artifact rejection, the function pop_artmwppth in the ERPLAB Toolbox was applied to all channels to discard epochs contaminated by residual blinks or other signals exceeding the designated amplitude thresholds (± 30 µV for horizontal EOG; ± 50 µV for vertical EOG and ± 80 µV for other channels). Data from participants with artifacts in more than 30% of trials were excluded from the subsequent ERP and behavioral analyses. The percentage of rejected trials ranged from 0.67% to 22.43%. For artifact correction, independent component analysis (ICA) was applied to all electrodes except for the mastoid electrodes and the EOG channels using the function pop_runica in the EEGLAB Toolbox. Then, the independent components (ICs) were classified into seven categories: brain, muscle, eye, heart, line noise, channel noise, and other sources using the ICLabel EEGLAB extension (Pion-Tonachini et al., 2019). The ICs were excluded if (1) the classification of each IC was eye, muscle, or channel noise with a high probability (>90%) and (2) the IC time course activity resembled a blink or vertical/horizontal eye movements or the power spectrum more closely resembled noise or muscle activity more than neural activity. For each participant, one to four ICs were excluded.

### Data Analysis

Averaged ERP waveforms were computed given a 1000 ms epoch starting 200 ms before (as baseline) and ending 800 ms after stimulus onset. For the exploration of the processing stages of the SNARC effect, the interaction between the SNARC and Stroop effects and between the SNARC and Simon effects for the P1, the N2, and P300 components were considered. The time window for each component was estimated using the collapsed localizers approach to avoid biased measurement (Chen et al., 2023; Luck & Gaspelin, 2017). Using this approach, the ERP waveforms from each condition were averaged. Then, the time windows were defined as the window between the onset and offset of each component in this averaged waveform. Precisely, the onset and offset latencies were measured as the 50% fractional peak latency (the latency at which the voltage reached 50% of the peak amplitude) (Kiesel et al., 2008). The P1 amplitude was calculated as the average amplitude at Fz between 121 and 195 ms after the onset of the target. The N2 amplitude was calculated as the average amplitude at FCz between 220 and 245 ms. The P300 amplitude was calculated as the average amplitude at Pz between 311 and 497 ms.

For statistical analysis, the mean amplitudes and the mean reaction times (RTs) of correct responses were considered dependent variables in a three-factor repeated measures analysis of variance (repeated-measures ANOVA, alpha = 0.05), including the 2 × 2 × 2 factors assessed in the SNARC, magnitude Stroop, and Simon effects. For the analysis of RTs, latencies faster than 200 ms and more than three standard deviations above the mean were excluded. Bonferroni correction was used for pairwise comparisons. Effect sizes are reported in terms of partial η^2^ and Cohen’s d. Meanwhile, the Bayes factor using the Jeffreys-Zellner-Siow (JZS) default prior was reported (Heck, 2019; Rouder et al., 2012). The Bayes factor quantified the likelihood of data fitting the H_1_ (alternative hypothesis) versus H_0_ (null hypothesis). The subscript of the *BF* refers to the models considered as the numerator and denominator for comparison. Accordingly, the Bayes factor for the alternative relative to the null model is denoted BF_10_, and BF_01_ denotes the null relative to the alternative model, i.e., 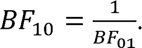 A BF_10_ greater than 30 indicates strong evidence in support of the alternative hypothesis. Conversely, a BF_10_ less than 1/30 suggests strong evidence in favor of the null hypothesis (Jeffreys, 1998; Wagenmakers et al., 2018).

All data were analyzed using R 4.1.2 (R Development Core Team, 2021), and any effect with more than one degree of freedom was adjusted for sphericity violations using the GreenhouseLGeisser correction. All *post hoc* analyses were Bonferroni-corrected. Pearson’s correlations were used to examine the relationship among measures.

## Results

### Reaction Times (RTs)

The results of repeated-measures ANOVA showed that the main effect of the SNARC was significant [*F*(1, 23) = 6.38, *p* = .019, 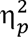 = .22, *BF_10_* > 30]. Moreover, an obvious RT advantage for the SNARC congruent condition was revealed (congruent: 486 ± 12 ms; incongruent: 505 ± 13 ms). Meanwhile, a significant main effect of the Stroop effect was observed [*F*(1, 23) = 20.14, *p* < .001, 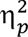 = .47, *BF_10_* > 30] and an obvious RT advantage for the Stroop congruent condition was revealed (congruent: 490 ± 12 ms; incongruent: 501 ± 11 ms). A significant main of the Simon effect was observed [*F*(1, 23) = 12.88, *p* = .002, 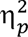 = .36, *BF_10_* > 30], with an obvious RT advantage for the Simon congruent condition (congruent: 493 ± 11 ms; incongruent: 499 ± 12 ms). Most importantly, both the interaction between SNARC and Stroop effects [*F*(1, 23) = 7.19, *p* = .013, 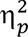 = .24, *BF_10_* = 1.01] and between the SNARC and Simon effects [*F*(1, 23) = 15.26, *p* < .001, 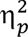 = .40, *BF_10_* > 30] were significant. Statistically, the SNARC effect is measured as the subtraction of the RT in the congruent condition from that in the incongruent condition. The SNARC effect in Stroop incongruent trials (15 ± 7 ms) was smaller than that in Stroop congruent trials (23 ± 8 ms) [*t*(23) = -2.71, *p* = .012, Cohen’s *d* = -.55, *BF_10_* = 4.04]. Similarly, the SNARC effect in Simon incongruent trials (26 ± 7 ms) was smaller than that in Simon congruent trials (13 ± 8 ms) [*t*(23) = -3.92, *p* < .001, Cohen’s *d* = -.8, *BF_10_* > 30]. However, there was no evidence of a two-way interaction between the Stroop and Simon effects [*F*(1, 23) = 0.88, *p* = .359, 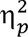 = .04, *BF_10_* = 0.05] or three-way interaction among the SNARC, Stroop and Simon effects [*F*(1, 23) = 1.61, *p* = .217, 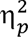 = .07, *BF_10_* = 0.11].

### P1

For the P1 component, only a significant interaction between the SNARC and Stroop effects was observed [*F*(1, 23) = 6.09, *p* = .021, 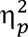 = .21, *BF_10_* > 30]. Here, the SNARC effect still refers to the result of subtracting the average amplitude within a particular time range in the congruent condition from that in the incongruent condition. *Post hoc* analysis showed that the size of the SNARC effect was greater in Stroop incongruent trials (-0.52 ± 0.23 µV) than in Stroop congruent trials (0.09 ± 0.25 µV) [*t*(23) = -2.25, *p* = .035, Cohen’s *d* = -.46, *BF_10_* = 1.75]. Furthermore, there was no evidence for the modulation of the SNARC effect by the Simon effect [*F*(1, 23) = 0.52, *p* = .476, 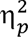 = .02, *BF_10_* < 0.03] or an interaction between the Stroop and Simon effects [*F*(1, 23) = 0.42, *p* = .521, 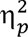 = .01, *BF_10_* = 0.89]. These findings indicated that only the congruency of the Stroop effect modulated the SNARC effect in the P1 component.

### N2

For the N2 component, the ANOVA revealed a significant main effect of the Stroop condition [*F*(1, 23) = 4.75, *p* = .040, 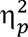 = .17, *BF_10_* = 1.09], with the N2 amplitude being significantly greater for the Stroop incongruent (-1.08 ± 0.79 µV) trials than for the Stroop congruent trials (-0.73 ± 0.77 µV). Meanwhile, a nonsignificant trend was found for the SNARC effect [*F*(1, 23) = 3.97, *p* = .058, 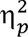 = .14, *BF_10_* = 12.18]. Most importantly, a significant interaction between the SNARC and Simon effects was observed [*F*(1, 23) = 6.42, *p* = .019, 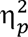 = .21, *BF_10_* = 5.37]. *Post hoc* analysis showed that the SNARC effect was greater in Simon congruent trials (-0.88 ± 0.30 µV) than in Simon incongruent trials (-0.20 ± 0.30 µV) [*t*(23) = -2.25, *p* = .019, Cohen’s *d* = -.52, *BF_10_* = 2.90]. Additionally, there was no evidence for the modulation of the SNARC effect by the Stroop effect [*F*(1, 23) = 0.35, *p* = .558, 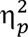 = .01, *BF_10_* = 0.23] or an interaction between the Stroop and Simon effects [*F*(1, 23) = 0.75, *p* = .395, 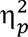 = .03, *BF_10_* = 3.44]. These findings indicated that only the congruency of the Simon effect modulated the SNARC effect for the N2 component.

### P300

For the P300 component, the ANOVA revealed a significant main effect of the SNARC condition [*F*(1, 23) = 4.78, *p* = .039, 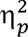 = .17, *BF_10_* > 30], with the P300 amplitude being significantly greater for the SNARC incongruent (10.04 ± 1.02 µV) trials than for the SNARC congruent trials (9.25 ± 0.89 µV). Most importantly, a significant interaction between the SNARC and Simon effects was observed [*F*(1, 23) = 8.56, *p* = .008, 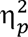 = .27, *BF_10_* = 11.41]. *Post hoc* analysis showed that the SNARC effect was greater in Simon incongruent trials (1.16 ± 0.37 µV) than in Simon congruent trials (0.42 ± 0.39 µV) [*t*(23) = 2.93, *p* = .008, Cohen’s *d* = .60, *BF_10_*= 6.11]. Additionally, there was no evidence for the modulation of the SNARC effect by the Stroop effect [*F*(1, 23) = 0.01, *p* = .947, 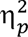 = .01, *BF_10_* = 0.11] or an interaction between the Stroop and Simon effects [*F*(1, 23) = 0.02, *p* = .883, 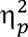 = .01, *BF_10_* = 4.67]. These findings indicated that only the congruency of the Simon effect modulated the SNARC effect for the P300 component.

## Discussion

To investigate the processing stage at which the SNARC effect occurs, the SNARC, Stroop (related to semantic representation), and Simon (related to response selection) effects were elicited in a magnitude comparison task and ERPs were measured. The behavioral results are in line with those reported in Nan et al. (2022), in which interactions of the SNARC and Stroop effects and of the SNARC and Simon effects were observed. More importantly, an interaction between the P1 components of the Stroop and SNARC effects was observed, which revealed that both the Stroop and SNARC effects involved a shared processing stage associated with early semantic representation. On the other hand, the interaction between the Simon and SNARC effects was observed for the conflict-associated N2 and P300 components, which further supported the involvement of both effects in the late response selection stage. In conclusion, the current findings support the proposition that the SNARC effect occurs flexibly at both semantic representation and response selection stages (Moro et al., 2018; Nan et al., 2022).

### The interaction between the Stroop and SNARC effects

The current behavioral results revealed a significant Stroop effect in which participants exhibited significant delays in RT when the semantic meaning of a task-irrelevant Chinese character was incongruent with the magnitude of the number. This indicates that the stimulus manipulation used to evoke the Stroop effect was effective. In the framework of the dimensional overlap (DO) theory, the Stroop effect in our study is caused by the overlap of the task-irrelevant (Chinese characters) and task-relevant (number) stimulus representation on semantic activation (Kornblum et al., 1990; Kornblum & Lee, 1995; Yan et al., 2021). Augustinova and Ferrand (2014) employed subtractive logic to demonstrate that semantic activation within the Stroop effect is automatic. The current study consistently found that the semantic processing of task-irrelevant Chinese characters remains automatic (Moors & De Houwer, 2006; Neely & Kahan, 2001)., even when the characters are partially masked by the target number.

Furthermore, an interaction between the Stroop and SNARC effects in both the RT and the P1 component was observed. The P1 component is regarded as the earliest endogenous visual ERP component linked to the sensory analysis of stimulus features (Paz-Caballero & García-Austt, 1992; Taylor, 2002). Holmes et al. (2009) employed an oddball paradigm in which within-category and between-category color deviants were equally salient in isolation. They observed enhanced and earlier P1 and N1 components exclusively for the between-category deviants, suggesting that these ERP components index early perceptual processes in color categorization. Notably, in the current study, the semantic representation within the Stroop effect was generated as a result of automatic magnitude (either large or small) processing (Moors & De Houwer, 2006; Neely & Kahan, 2001). Following the logic of the AFM, the interaction in the P1 component might indicate that the SNARC effect involves processing of the magnitude representation, which is also involved in the Stroop effect.

The debate regarding whether the spatial–numerical associations are derived from the magnitude or ordinal representation has been long-standing (Casasanto & Pitt, 2019; Pellegrino et al., 2019; Pinto et al., 2021). According to the two-stage processing model, the association between numbers and space results from the specific manner of encoding magnitude representation (Dehaene et al., 1993; Gevers et al., 2006). Van Dijck and Fias (2011) challenged magnitude representation by inserting a parity judgment task during the maintenance interval of a verbal working memory (WM) task. Participants were instructed to solely judge the parity of the numbers they maintained in a random order within WM. The results showed associations between position and spatial representation, rather than between magnitude and spatial representation, with items from the beginning/end of a verbal WM sequence showing faster and more accurate responses when using the left/right hand. A working memory account for that phenomenon was proposed, which underscores the crucial role of ordinal WM representation, rather than magnitude representation, in spatial–numerical association (Gevers et al., 2003; Majerus & Oberauer, 2020; van Dijck et al., 2009). Recently, Koch et al. (2023) cleverly selected number sets with a critical number that dissociates ordinal position and magnitude to investigate the individual contributions of magnitude and ordinal representation to spatial–numerical association. Their results contradict those obtained by considering WM, as the association can rely on either magnitude or ordinality (Prpic et al., 2016). Even though in the current experiment the ordinal representation was not accounted for, the observation of the interference from the magnitude Stroop effect suggests that the inclusion of magnitude is crucial when exploring the underlying mechanisms of the SNARC effect.

### The interaction between the Simon and SNARC effects

The Simon effect induced in the current experiment showed that participants were remarkably faster if the orientation of the number spatially corresponded to the correct response. The interaction between the Simon and SNARC effects is still controversial (Gevers et al., 2005; Mapelli et al., 2003). This controversy may be partially attributed to the use of different types of Simon effects, such as visuomotor Simon or cognitive Simon effects (Xiang et al., 2022; Yan et al., 2023). The visuomotor Simon effect is derived from the interference of exogenous spatial representation (i.e., the location of stimuli). This interference is computed by the visuomotor pathways, and its effect size decreases until it disappears at the slowest RTs (Pellicano et al., 2009; Pratte et al., 2010). The cognitive Simon effect, as introduced in the current experiment, is derived from the conflict between cognitive representations, and its size is enhanced with increasing RT (Ansorge, 2003; Yan et al., 2021). Previous research using RT distribution analysis (De Jong et al., 1994) and drift diffusion model analysis (Ratcliff et al., 2016) has provided evidence to support the idea that the SNARC effect is more similar to the cognitive Simon effect than to the visuomotor Simon effect (Yan et al., 2023).

In line with previous studies, the current study findings support the similarity of the cognitive Simon and SNARC effects and further provides data reflecting the temporal dynamic characteristic of this similarity. Interactions between these two effects was observed for both the N2 and P300 components. The N2 and P300 components are considered neural markers of different subprocesses of cognitive control (Groom & Cragg, 2015; Li et al., 2015; Wang et al., 2014). For example, Groom & Cragg (2015) conducted a hybrid go/no-go task and observed that while amplitudes of both N2 and P300 components are enhanced during the conflict, N2 shows a sensitivity to the degree of conflict, whereas P300 exhibits a sensitivity toward the required degree of response inhibition. Consistently, the current ERP results showed that the amplitude of the N2 component was enhanced in Stroop incongruent trials. However, the SNARC and Simon effects are somewhat different. The N2 amplitude was enhanced in the SNARC incongruent trials. However, the extent of the enhancement was reduced in the trials in which the Simon effect occurred. It seems that the processing resulting in the Simon effect competes for cognitive resources involved in monitoring the SNARC conflict (Botvinick et al., 2001). This modulation of the SNARC effect by the Simon effect is also manifested in the P300 component. The P300 amplitude was enhanced in the SNARC incongruent trials, while the extent of this enhancement was further amplified in the trials in which the Simon effect occurred. The findings suggest that the conflict resolution mechanism of the SNARC and Simon effects shared conflict monitoring and response inhibition processing, where the two compete for limited cognitive resources (Gevers et al., 2005; Keus & Schwarz, 2005; Yan et al., 2021).

### The SNARC effect in the framework of the two-stage processing model

Notably, Yan et al. (2022) and Xiang et al. (2022) proposed a two-stage processing model to explain the occurrence of the SNARC effect. This two-stage processing model refines our understanding of the spatial-numerical association by subdividing the SNARC effect mechanistically into two process stages, the spatial representation of the magnitude and the spatial representation of the response selection, which are similar to the semantic-representation stage and response-selection stage, respectively. The current study verified this model with ERP evidence. However, this dual-stage proposition seems inconsistent with some research findings in which the SNARC effect is manifested as a single-stage effect (Keus & Schwarz, 2005; Mapelli et al., 2003; Yan et al., 2021). The inconsistency might be attributed to the difference in introduced interference factors since the two-stage processing model proposes that different interference factors acting on either of the two stages might be the core reason for the stage flexibility of the SNARC effect (Xiang et al., 2022; Yan et al., 2022). Xiang et al. (2022) confirmed the dominant influence of the magnitude-relevance of interference information on the SNARC effect. Notably, in the current study, the interaction of the magnitude Stroop and SNARC effects was exclusively observed for the P1 component but absent for the later components. This raises the possibility that the interaction between the SNARC and Stroop effects observed in previous studies (Nan et al., 2022) is mainly due to a shared magnitude representation; in contrast, when the interference (e.g., parity information) is magnitude-independent (Yan et al., 2021), it does not influence the SNARC effect, showing that the SNARC effect manifests as a single-stage effect.

Numerous researchers have attempted to explain the SNARC effect from various perspectives (Dehaene et al., 1993; Gevers et al., 2006; Proctor & Cho, 2006; van Dijck & Fias, 2011). However, our understanding is still not complete since the relationship between the numbers’ spatial representation and the response has been overlooked. How does spatial representation, activated by the perception of numbers, conflict with response processing during response selection? The second stage of the two-stage model, namely, the spatial representation of the response selection, provides a description of this process. The model employs three levels (the input, hidden, and output levels). In the output level, where the response selection takes place, the automatically encoded spatial representation of the number and task-critical semantic representation enhance their corresponding spatial responses. If the response corresponding to the spatial representation is incongruent with that of the semantic representation, cognitive control will inhibit the spatial representation and result in a delay in response, known as the SNARC effect.

The interaction between the SNARC and cognitive Simon effects, which indicates that both effects shared conflict monitoring and response inhibition processing, render the description of the two-stage processing model meaningful. According to the theory of event coding (TEC), both stimuli and response representations are represented in a common coding system using feature codes (Hommel et al., 2001). The stimulus representations are compound codes representing the stimulus location and other attributes. Activation of the feature codes of a left stimulus spreads to automatically activate those of the left response (Hommel, 1993). Likewise, the response representation is compound of encodings of the motor patterns and the resulting perceptual consequences. A left-hand response, because it occurs on the left side of space, is also encoded as “left” spatial features. The Simon effect arises due to the sharing of the spatial features between stimulus and response codes in the framework of TEC (Hommel, 2011; Van der Lubbe & Abrahamse, 2011). Considering that the Simon and SNARC effects might share a conflict resolution mechanism, the spatial representation of the number would similarly impact the response selection, as it shares spatial features with the response representation in the response selection stage.

### Limitation

Notably, Dehaene et al. (1993) initially discovered the SNARC effect in a parity judgment task. Previous studies confirmed the heterogeneity of the cognitive resources invoked in the parity judgment task and magnitude comparison task (Deng et al., 2017, 2018). Deng et al. (2017) conducted a series of experiments and claimed that the strength of the SNARC effect fluctuated over time in parity tasks but increased with time in magnitude tasks and remained more stable. In future studies, the relationship between the processing stages of the SNARC effect and the task set needs to be considered.

## Conclusion

Derived from high temporal resolution EEG recordings, the current findings support the two-stage processing model, ultimately indicating that the occurrence of the SNARC effect may involve both the semantic representation and response selection stages.

## Acknowledgments

We thank all people involved in the data acquisition process. We would like to acknowledge all members of the Center for Brain and Cognitive Sciences for their continued support.

## Compliance with ethical standards

### Funding

This work was supported by grants from the Youth Project of Basic and Applied Basic Research Fund of Guangdong Province - Regional Joint Fund (No. 2021A1515110452).

### Ethical approval

This study was approved by the Institutional Review Board of the Educational School, Guangzhou University. All procedures in studies involving human participants were performed in accordance with the ethical standards of the institutional and national research committees and with the 1964 Declaration of Helsinki and its later amendments or comparable ethical standards.

### Informed consent

Informed consent was obtained from all individual participants included in the study.

## Author Contributions

**Weibin Yang:** Investigation; validation; formal analysis; visualization; project administration; writing – original draft. **Lizhu Yan:** Conceptualization; methodology; software; investigation; data curation. **Xiangxin Rui:** Methodology; software; data curation; project administration. **Weizhi Nan:** Conceptualization; funding acquisition; investigation; methodology; project administration; resources; supervision; writing – review and editing.

## Data Accessibility

The anonymized data, stimuli, and preprocessing/analysis details are available upon request.

